# Integrative Transcriptomic and Machine Learning Approaches to decipher Mitochondrial Gene Regulation in severe *Plasmodium vivax* Malaria

**DOI:** 10.1101/2025.07.01.662590

**Authors:** Priyanka Roy, Yash Aggarwal, Sanjay Kumar Kochar, Dhanpat Kumar Kochar, Ashis Das

## Abstract

Mitochondria in *Plasmodium vivax* are functionally vital despite possessing a highly reduced genome and differing substantially from the human organelle. Beyond their classical role in energy production, they dynamically coordinate processes like pyrimidine biosynthesis and heme metabolism, adapting their functions across the intra-erythrocytic development cycle (IDC). Their unique architecture and stage-specific roles enable the parasite to fine-tune mitochondrial gene expression, involving both protein-coding sense transcripts and long non-coding natural antisense transcripts (NATs). This study unveils an unprecedented regulatory complexity by integrating transcriptomic profiling with advanced machine learning to decode the role of mitochondrial sense and natural antisense transcripts (NATs) in severe *P. vivax* malaria. We reveal distinct, clinically relevant expression signatures, where NATs emerge not as transcriptional by-products but as potent regulators tightly linked to mitochondrial pathways and translational machinery. This dual-layered transcriptomic landscape reflects an intricate molecular strategy by which the parasite fine-tunes mitochondrial function to survive under severe disease conditions. Importantly, while these findings illuminate novel regulatory mechanisms and position mitochondrial NATs as promising targets for antimalarial drug development, they represent preliminary insights derived from a limited clinical cohort and should not be interpreted as definitive clinical indicators. Validation in larger and diverse patient populations is essential to confirm their broader biological and clinical relevance. However, these results serve as indicators for potential innovative therapeutic interventions aimed at disrupting parasite bioenergetics and regulatory networks.

## Introduction

Malaria, a significant global health challenge, is caused by protozoan parasites of the genus *Plasmodium*, with *Plasmodium vivax* being one of the most widespread and prevalent species. According to the World Health Organization (WHO) 2024 report, an estimated 263 million malaria cases were recorded globally in 2023. Strikingly, *Plasmodium vivax* represented approximately 48.2% of malaria infections in the South-East Asia region, with India emerging as a major contributor, accounting for nearly 25% of the global *P. vivax* burden [1], highlighting its substantial burden on public health. Historically, *P. vivax* was thought to cause benign tertian malaria, however many studies have reported that it causes significant severity & morbidity, where changes in gene expression plays a major role on the underlying mechanisms contributing to this shift. Transcriptional analyses have shown that variations in gene expression profiles can affect the parasite’s ability to invade erythrocytes and modulate host immune responses [2,3].

Most eukaryotic genomes produce non-coding RNAs (ncRNAs), which integrate complex suites of gene activity and contribute to the complexity of gene regulation [4]. The lncRNAs are a subclass of non-coding RNAs (ncRNAs) that are longer than 200nt and are typically distinguished by their inability to code for proteins. They can be categorized as intergenic, intronic, sense lncRNAs, transcribed from the sense strand of protein-coding genes, and Natural Antisense Transcripts (NATs), transcribed from the antisense strand of protein-coding genes, based on the genomic area of transcription [4]. The lncRNAs can modulate gene expression at numerous levels, including transcription, post-transcription, RNA turnover and protein translation [6,7].

The presence of ncRNA in *Plasmodium* has the potential to regulate key events in the complex life cycle and modulate the virulence ability of the parasite, by controlling transcriptional and post-transcriptional processes [8]. Patankar et al.,2001 & Militello et al., 2005 reported the presence of antisense transcripts in *Plasmodium* parasite and their role in gene regulation [9,10]. Additionally, the presence of antisense transcripts in gametocytes and ookinetes implies that these antisense RNAs could be essential for the control of gene expression and parasite development [11]. Studies from our laboratory have showed the presence of NATs using a custom designed strand specific whole genome microarray in *P. vivax* from patients showing complicated and uncomplicated disease manifestations [12].

In *Plasmodium* the role of the mitochondria is particularly fascinating because its function varies depending on the stage of the parasite’s lifecycle. It has a minimalistic genome encoding only three proteins cytc oxidase subunit 1 (cox1) and subunit 3 (cox3), members of the cytc oxidase complex (complex IV), and cytb (cob), a member of cytochrome bc1 complex. Despite the reduced genome, *Plasmodium* mitochondria play crucial roles in metabolism, including pyrimidine biosynthesis and electron transport chain in the asexual blood stages [13, 14]. Apart from this, there is a functional citric acid cycle within the mitochondrion which is responsible for the de novo biosynthesis of heme. Despite the widespread haemoglobin breakdown in the digestive vacuole, heme production in *Plasmodium* is divided between the mitochondrion, the apicoplast, and the cytosol [13,14,15,16].

Despite mitochondria having their own genome with transcriptional and translational machinery, majority of the proteins found in mitochondria are encoded in the nucleus due to a huge endosymbiotic gene transfer from the mitochondrial to the nuclear genome [17]. There are possible energy requirement differences between the asexual intra-erythrocytic developmental stages and the sexual stages such as gametocytes. In asexual stages, no cristae are visible in the mitochondrial membrane while in the mitochondria of gametocytes, crista-like structures accumulate in the respiratory chain complexes [18, 19]. Another interesting observation remains that there is a significantly lower level of Complex V components in the asexual stages, compared to the sexual stages of the parasites [19].

Even though the *Plasmodium* mitochondria has limited number of metabolic process & pathways, mitochondrial gene products and their expression are functionally vital, with stage-specific remodelling evident between asexual and gametocyte stages, highlighting their fundamental contribution to parasite viability and transmission potential. A wide range of drugs are designed to target mitochondrial pathways, but the emergence of resistance has become a significant concern during their development. The relaxed selective pressure for mitochondrial electron transport chain function during the blood stage, along with mitochondrial genome polyploidy seems to accelerate the emergence of atovaquone-resistance mutations [20].

Based on the data in this study, this is the first comprehensive evidence of mitochondrial transcripts from *P. vivax* clinical isolates with differing disease manifestations showing transcripts in both sense and NATs orientation. Through integrative transcriptomic profiling and machine learning models, we uncovered key insights into mitochondrial gene regulation mediated by NATs, particularly in the context of severe disease manifestation. Notably, we observed distinct differential expression patterns of NATs across clinical conditions, with a strong association to mitochondrial pathways and the translational machinery. These findings suggest that NATs are not random transcriptional by-products but potentially function as regulatory elements that fine-tune mitochondrial gene expression during pathogenesis. Altogether, in correlation with the gene expression signatures, the mitochondrial NATs in *Plasmodium* may provide alternative targets for antimalarial drug development and disease intervention strategies.

## Materials & Methods

### Curating gene list with potential mitochondrial localization

To employ a systematic approach to compile a comprehensive list of *P. vivax* mitochondrial genes, a multi-layered bioinformatic strategy was employed, integrating web-based prediction tools with functional annotations. Given the absence of a universally conserved mitochondrial localization signal in *Plasmodium* species and the limitations of individual prediction tools each trained on distinct datasets and algorithms multiple complementary platforms were utilized, following the recommendations of Martelli et al., 2021 [21]. The four primary tools included: TargetP, which identifies mitochondrial transit peptides (mTPs) characterized by enrichment of arginine, leucine, and serine residues critical for matrix targeting [22]; Cello2GO, which leverages BLAST-based homology to assign subcellular localization [23]; MitoFates, which predicts mitochondrial pre-sequences using classical targeting features and motif analyses [24]; and Mitoprot II, a tool that employs discriminant analysis of amino acid composition, charge, hydrophobicity, and amphiphilic potential to identify mitochondrially imported proteins [25]. Additionally, the analysis was strengthened by incorporating genes annotated with mitochondrial targeting domains in PlasmoDB [26], alongside orthologous genes that have been experimentally validated as part of the *Plasmodium falciparum* mitochondrial proteome [27–30]. This combined approach ensured the inclusion of both predicted and empirically confirmed mitochondrial candidates. To ensure prediction robustness, a stringent cutoff score of >0.6 was applied for each tool, and duplicate predictions across different platforms were consolidated. This filtering yielded a final non-redundant list of 617 unique nuclear-encoded, mitochondria-targeted candidates in *P. vivax*.

### Blood sample collection & processing

Venous blood samples were obtained from patients diagnosed with *Plasmodium vivax* infection at S.P. Medical College, Bikaner, India. To enrich for infected erythrocytes, the collected blood was processed using a Histopaque density gradient. All patient samples were collected by a dedicated team of clinicians under ethical approval, with informed consent obtained in accordance with hospital guidelines. The presence of *P. vivax* infection was initially confirmed by slide microscopy and rapid diagnostic tests (RDTs), followed by PCR-based validation to ensure mono-infection [31–32]. The patient’s diagnosis was confirmed as hepatic dysfunction as per WHO guidelines [33] by the team of clinicians. Additionally, no samples processed showed any co-morbidities. Detailed clinical characteristics of the enrolled patients are presented in Table S1. For transcriptomic profiling, total RNA from each sample was individually hybridized onto custom-designed 8×60K single-color microarrays.

### Microarray Hybridization

Total RNA was extracted from patient blood samples following established protocols [12]. The concentration and purity of RNA were assessed using a NanoDrop 2000 spectrophotometer (Thermo Scientific) and Qubit fluorometric assay (Thermo Scientific, USA). RNA integrity was evaluated on an Agilent TapeStation system to ensure high-quality input for downstream analysis. Microarray labeling and hybridization were carried out at the Agilent-certified facility of Genotypic Technology Pvt. Ltd., Bengaluru, India. RNA samples were processed using the Agilent Quick-Amp Labeling Kit (p/n 5190-0442). In this process, total RNA was reverse transcribed at 40°C using an oligo-dT primer linked to a T7 polymerase promoter to synthesize first-strand cDNA. This was followed by second-strand synthesis to generate double-stranded cDNA, which served as a template for in vitro transcription. During in vitro transcription, cyanine-3 (Cy3) labeled cRNA was produced using Agilent Cy3 CTP. All enzymatic reactions, including cDNA synthesis and transcription, were performed at 40°C. The labeled cRNA was purified using Qiagen RNeasy columns (Cat. No. 74106), and quality metrics—yield and specific activity—were confirmed using the NanoDrop ND-1000. Prior to hybridization, labeled cRNA was fragmented at 60°C and applied to Agilent Custom *P. vivax* 8×60K Gene Expression Microarrays. Hybridization was carried out using the Agilent Gene Expression Hybridization Kit (Part No. 5190-0404) in SureHyb chambers at 65°C for 16 hours. Post-hybridization washes were performed using Agilent Gene Expression Wash Buffers (Part No. 5188-5327). Microarray slides were scanned with the Agilent Microarray Scanner (Model G2600D), and raw data were extracted using Agilent Feature Extraction software from images for further analysis.

### Microarray design

A custom 60K sense-antisense microarray was meticulously designed to enable comprehensive transcriptomic profiling of *Plasmodium vivax*, with a particular emphasis on mitochondrial gene regulation. The array was constructed using genomic annotations from the *P. vivax* Sal-1 reference genome PlasmoDB v47 (www.plasmodb.org), incorporating both nuclear and mitochondrial open reading frames (ORFs). Each ORF was represented by five distinct oligonucleotide probes (60-132 mers) in both sense and antisense orientations. Additionally, probes targeting plasmodial mitochondrial sequences were cross-verified against the *Homo sapiens* mitochondrial genome and the probes for nuclear encoded transcripts were also cross-verified against the *Homo sapiens* nuclear genome; probes with significant matches were removed, any cross-hybridizing probes were also excluded. The array was fabricated on the Agilent microarray platform, ensuring optimal hybridization performance and data quality. Probes from our previously validated 15K and 244K microarray [3,12] were selectively integrated into the current 60K design to enhance coverage and reliability. In this study, we specifically report transcripts, detected through hybridization, that are either encoded by the *P. vivax* mitochondrial genome or are predicted to be mitochondrial-targeted.

### Microarray Data Analysis

Feature-extracted raw image data, filtered for signals at least two-fold above background intensity, were processed using Agilent GeneSpring GX Software. Data normalization was performed using the 75th percentile shift method to ensure consistency across samples and reduce technical variability. To enhance strict stringency, only probes detecting transcripts in at least 60% of samples were retained for further analysis. Differential gene expression was assessed by comparing complicated samples to uncomplicated samples. Genes showing a fold change ≥1.5 were considered biologically relevant, and their statistical significance was determined using a student t-test (p ≤ 0.05), with multiple testing correction applied via the Benjamini-Hochberg false discovery rate (FDR) implemented through the limma and genefilter R packages in Bioconductor.

### Machine Learning

The differentially expressed genes were statistically ranked by adjusted p-values and used to construct training datasets for machine learning. We use several indexes, including F1-Score, Matthews Correlation Coefficient (MCC), and Area Under the Curve (AUC), to evaluate the performance of models, each with its specific formula, significance, and application, particularly in scenarios with imbalanced data (non-uniform between uncomplicated and complicated samples).

The metric of F1-Score is the harmonic mean of precision and recall, calculated as

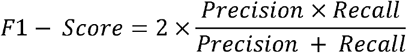

F1-Score, the harmonic mean of precision and recall, provides a balanced assessment of a model’s ability to identify both true positives and true negatives. A score close to 1 indicates strong performance, particularly vital in datasets with unequal class distributions.

MCC is calculated using the formula:

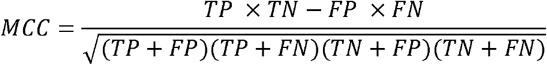

Where TP, TN, FP, and FN represent true positives, true negatives, false positives, and false negatives, respectively. It offers a comprehensive evaluation that considers all elements of the matrix (TP, TN, FP, FN). It ranges from -1 (inverse prediction) to +1 (perfect prediction). MCC is especially useful in imbalanced datasets for its balanced insight.

AUC (Area Under the Curve) quantifies a model’s ability to distinguish between classes, independent of class imbalance, AUC ranges from 0 to 1.

Unlike simple accuracy, these metrics offer nuanced insights, ensuring the model’s effectiveness is not masked by skewed class ratios. A high F1-Score, MCC, and AUC collectively indicate the model’s strength in accurately predicting both complicated and uncomplicated cases, providing a holistic assessment of its predictive capabilities across different class distributions.

To systematically evaluate predictive performance and feature stability, the differentially expressed genes were ranked by adjusted p-values and progressively reduced in subsets from top 92, 80, 70, 60, 50, 40, 30, and 20 transcripts for sense transcripts, and 77, 70, 60, 50, 40, 30, and 20 transcripts for NATs. This stepwise reduction enabled us to examine how the number of features influenced model robustness and interpretability. Our machine learning models, trained on these transcriptomic signatures, were further interrogated to uncover potential regulatory transcripts in sense & NATs that may act as drivers involved in the modulation of mitochondrial gene expression. The resulting models not only demonstrated high predictive performance but also guided downstream analyses to explore enriched pathways and biological functions. This integrative approach helps direct, preliminary studies, for identifying potential molecules and improving clinical stratification of malaria patients.

We employed three machine learning algorithms, including XGBoost [34] with parameters n_estimators=100,max_depth=2,learning_rate=0.05,subsample=0.8,colsample_bytree=0.8,re g_alpha=0.1,reg_lambda=1, random forest [35] with parameters for max_depth = 3, min_samples_split = 5, n_estimators = 100, logistic regression [36] with parameters max_iter = 1000 to test and predict key regulators of disease severity in case of mitochondrial transcripts from sense & NATs. The Leave-One-Out (LOOCV) cross-validation technique was utilized [37].

Due to the limited number of samples, the dataset was not further split into independent training and testing sets, as this would have resulted in very small partitions. Instead, we employed Leave-One-Out Cross-Validation (LOOCV) to address critical challenges in evaluating machine learning models trained on limited and high-dimensional biological data. For imbalanced datasets, standard cross-validation may create folds with zero minority-class samples, skewing performance metrics. Small datasets with high dimensionality are prone to overfitting. LOOCV’s exhaustive resampling helps detect overfitting by testing the model’s consistency across all possible training configurations. In standard k-fold cross-validation, the dataset is partitioned into k subsets, with each subset serving as the test set once. LOOCV represents the extreme case of k-fold CV, where k equals the total number of samples. LOOCV guarantees each test set contains exactly one sample, preserving class representation and providing a more reliable estimate of model sensitivity/specificity.

For each iteration of LOOCV, N-1 samples are used to train the model while the remaining single sample serves as the test set. This process is repeated N times, ensuring every sample is used as the test set exactly once.

Mathematically, the LOOCV estimate of the mean squared error (MSE) is given by:

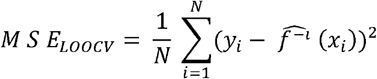

where N denotes the number of iterations, and the term in the brackets denotes the model trained on all samples except the i-th observation.

LOOCV helps to ensure no data wastage, which is ideal for precious samples, and provides stability by removing randomness from fold generation, which is crucial for publishing reproducible results. Computational cost for employing this technique increases linearly with the sample size, making it prohibitive for large datasets. In our study, LOOCV ensured rigorous validation without sacrificing statistical robustness.

To gain deeper insights into model interpretability, we employed SHAP (SHapley Additive exPlanations, v0.47.2) to unravel the contribution of individual features to the prediction outcome. By computing Shapley values, we quantified the impact of each feature on the model’s decision-making process, enabling a precise, feature-level understanding of predictive influence [38]. Genes were ranked by mean absolute SHAP values across LOOCV folds. To strengthen prioritization, we also examined stability across models and folds, and cross-validated biological relevance using gene set enrichment analysis. Only genes meeting both statistical and biological criteria were emphasized.

### Permutation Testing for Model Validation

To rigorously assess whether our machine learning models captured genuine biological signal rather than chance correlations, we employed permutation testing as a negative control. In this approach, the class labels (uncomplicated vs complicated) were randomly shuffled while preserving the gene expression structure, thereby destroying any true association between features and labels. The models (Random Forest, Logistic Regression, XGBoost) were retrained and evaluated on these permuted datasets across 200 iterations, and performance metrics were recalculated. The distribution of scores obtained from permuted datasets represents the null distribution the level of performance expected if the models were learning noise only. By comparing the observed performance on the real data to this null distribution, we calculated empirical p-values, defined as the proportion of permuted scores greater than or equal to the observed score. This approach directly tests whether the observed model performance is statistically greater than chance.

### Gene set enrichment and pathway analysis

To uncover the biological meaning of the machine learning prioritized signatures, we carried out gene set enrichment analysis (GSEA) on the final optimized subsets of transcripts i.e. 80 sense transcripts and 70 NATs that achieved the best balance between predictive accuracy and interpretability. GSEA included Gene Ontology (GO) term prediction and pathway enrichment using the PlasmoDB resource [26], which provided a user-friendly interface for exploring parasite biology. Analyses were performed with a significance threshold of p = 0.05, and pathway information was obtained from KEGG [39] and MetaCyc [40]. Focusing enrichment only on the optimized transcript sets avoided the noise and overfitting associated with very large or arbitrary gene lists, ensuring that the functional interpretations reflected the most robust biological signals. This strategy allowed us to directly link model-selected features to meaningful pathways relevant to regulatory network in the mitochondria.

## Results

Our previous studies have also documented *P. vivax* infections presenting with severe complications such as cerebral malaria, acute respiratory distress syndrome, jaundice, acute renal failure, etc., in adult patients [12]. Building on these findings, the current study focuses on 14 adult cases of *P. vivax* malaria, encompassing both complicated and uncomplicated manifestations, classified in accordance with WHO guidelines [34]. Detailed clinical characteristics of these patients are provided in Table S1.

### Mitochondrial associated genes

A curated list of 636 mitochondrial-associated genes was compiled, encompassing 19 genes encoded by the mitochondrial genome and 617 nuclear-encoded genes predicted to target the mitochondria. These predictions were derived using a suite of tools including TargetP [22], Cello2GO [23], MitoFate [24], MitoprotII [25], PlasmoDB [26] and orthologous genes that have been experimentally validated as part of the *Plasmodium falciparum* mitochondrial proteome [27–30]. To ensure the robustness of the gene selection, a stringent criterion was applied with a cutoff score >0.6 for each tool. The comprehensive list provided in (Supplementary Data-2) comprises computationally predicted nuclear-encoded mitochondrial-targeted transcripts along with core mitochondrial transcripts, forming the basis for the transcriptomic analyses in this study

### Differential gene expression analysis in clinical isolates

To uncover the differential gene expression patterns between complicated and uncomplicated *P. vivax* malaria, we normalized microarray data from 14 clinical samples using the 75th percentile shift method. Differential expression analysis was performed using a fold change threshold of ≥1.5 and a significance level of p ≤ 0.05, with correction for multiple testing via the Benjamini-Hochberg false discovery rate method. Expression fold changes were calculated as the average signal intensity across complicated malaria samples. To ensure biological relevance, only probes mapping to *P. vivax* Sal-I transcript sequences and detected in at least 9 out of 14 samples were considered for further analysis. From a total of 636 mitochondrial transcripts, a total of 626 genes were analysed for sense transcripts and 581 genes for NATs (Supplementary Data-3). Out of those a total of 92 and 77 genes were differentially expressed for sense and NATs respectively, in complicated *P. vivax* malaria samples as compared to the uncomplicated *P. vivax* malaria samples (Supplementary Data-3). Among the 92 sense transcripts, 66 genes (71.7%) were upregulated, while 26 genes (28.3%) were downregulated and out of 77 differentially expressed NATs, 27 genes (35.1%) were upregulated, and a notable 50 genes (64.9%) were downregulated (Fig-1).

**Figure 1.**
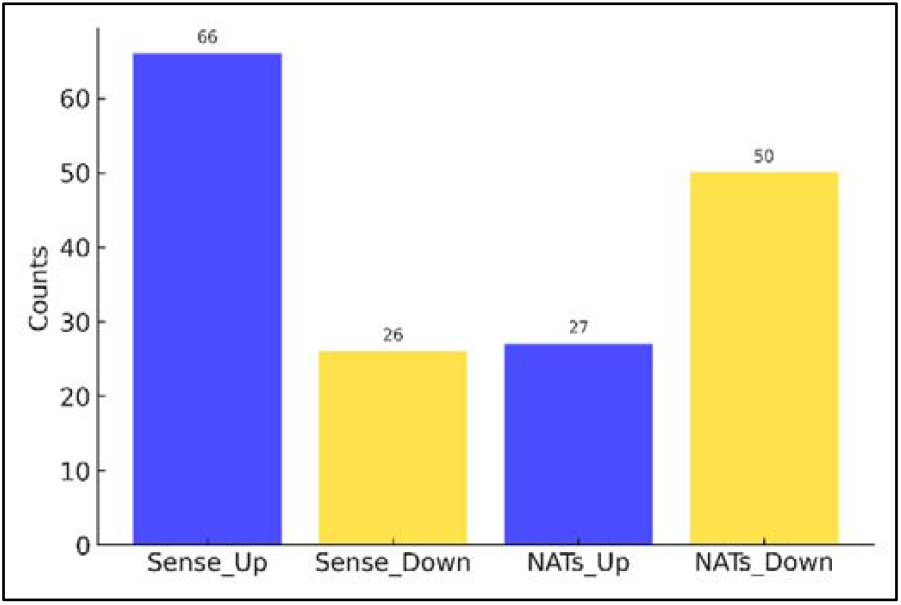
Bar plot illustrating the distribution of differentially expressed Sense and NATs transcripts

### Evaluating Machine Learning Sensitivity Across Varying Thresholds of Significant Gene Expression

To explore the predictive potential of mitochondrial transcripts, our initial analysis focused on evaluating features derived from statistically significant differentially expressed genes (DEGs) in both sense and NATs. These features were assessed for their utility in machine learning aimed at identifying key markers or genes in mitochondrial regulation. From the differential expression analysis, 92 sense and 77 NATs were shortlisted based on their adjusted p-values (Supplementary Data-4), and subsequently prioritized for downstream predictive modelling and pathway enrichment. To gauge predictive strength, we systematically evaluated subsets of these DEGs ranging from the top 92 to 20 for sense transcripts, and from 77 to 20 for NATs (Supplementary Data-4), and then used three different machine learning algorithms. Remarkably, performance metrics including accuracy, F1-score, and AUC consistently approached or exceeded 90% for subsets ranging from the top 80 to 20 in sense, and 70 to 20 in NATs. These findings underscore the robustness of these gene sets in predictive modelling and highlight their potential biological relevance. To further assess the stability of these models, we computed standard deviations across LOOCV iterations and repeated the process with random shuffling both for sense & NATs (Table 1, 2). Based on the comparative evaluation, the top 80 genes in sense and top 70 in NATs offered the most reliable prediction performance across all algorithms. Accordingly, these subsets were selected for further gene set enrichment analysis to enable a deeper, systems-level understanding of the mitochondrial regulatory landscape potentially governed by these transcripts.

**Table 1.** Machine learning results of the top 92, 80, 70, 60, 50, 40, 30, 20 significantly expressed genes, and randomly selected 40 genes in Sense.

|  |  |  |  |  |  |  |  |  |  |  |  |
| --- | --- | --- | --- | --- | --- | --- | --- | --- | --- | --- | --- |
| Model | Metric | 92 genes | 80 genes | 70 genes | 60 genes | 50 genes | 40 genes | 30 genes | 20 genes | Random 40 genes | Standard Deviation |
| Random Forest | Accuracy | 0.867 | 0.933 | 0.933 | 0.933 | 0.933 | 0.933 | 0.933 | 0.933 | 0.800 | 0.269 |
|  | Precision | 0.889 | 0.900 | 0.900 | 0.900 | 0.900 | 0.900 | 0.900 | 0.900 | 0.750 | 0.508 |
|  | Sensitivity | 0.889 | 1.000 | 1.000 | 1.000 | 1.000 | 1.000 | 1.000 | 1.000 | 1.000 | 0.508 |
|  | Specificity | 0.833 | 0.833 | 0.833 | 0.833 | 0.833 | 0.833 | 0.833 | 0.833 | 0.500 | 0.487 |
|  | F1 Score | 0.889 | 0.947 | 0.947 | 0.947 | 0.947 | 0.947 | 0.947 | 0.947 | 0.857 | 0.508 |
|  | MCC | 0.722 | 0.866 | 0.866 | 0.866 | 0.866 | 0.866 | 0.866 | 0.866 | 0.548 | 0 |
|  | AUC | 0.981 | 0.981 | 0.981 | 0.981 | 1 | 0.963 | 1 | 1 | 0.954 | 0.022 |
| Logistic Regression | Accuracy | 0.933 | 0.933 | 0.933 | 0.933 | 0.933 | 0.933 | 0.933 | 0.933 | 0.6 | 0.032 |
|  | Precision | 0.9 | 0.9 | 0.9 | 0.9 | 0.9 | 0.9 | 0.9 | 0.9 | 0.667 | 0.507 |
|  | Sensitivity | 1 | 1 | 1 | 1 | 1 | 1 | 1 | 1 | 0.667 | 0.507 |
|  | Specificity | 0.833 | 0.833 | 0.833 | 0.833 | 0.833 | 0.833 | 0.833 | 0.833 | 0.5 | 0.504 |
|  | F1 Score | 0.947 | 0.947 | 0.947 | 0.947 | 0.947 | 0.947 | 0.947 | 0.947 | 0.667 | 0.507 |
|  | MCC | 0.866 | 0.866 | 0.866 | 0.866 | 0.866 | 0.866 | 0.866 | 0.866 | 0.167 | 0 |
|  | AUC | 0.926 | 0.926 | 0.926 | 0.944 | 0.944 | 0.963 | 0.981 | 1 | 0.759 | 0 |
| XGBoost | Accuracy | 0.8 | 0.8 | 0.867 | 0.867 | 0.8 | 0.8 | 0.8 | 0.8 | 0.867 | 0.433 |
|  | Precision | 0.692 | 0.692 | 0.692 | 0.692 | 0.692 | 0.692 | 0.692 | 0.692 | 0.818 | 0.507 |
|  | Sensitivity | 1 | 1 | 1 | 1 | 1 | 1 | 1 | 1 | 1 | 0.507 |
|  | Specificity | 0.333 | 0.333 | 0.333 | 0.333 | 0.333 | 0.333 | 0.333 | 0.333 | 0.667 | 0.380 |
|  | F1 Score | 0.818 | 0.818 | 0.818 | 0.818 | 0.818 | 0.818 | 0.818 | 0.818 | 0.9 | 0.507 |
|  | MCC | 0.48 | 0.48 | 0.48 | 0.48 | 0.48 | 0.48 | 0.48 | 0.48 | 0.739 | 0 |
|  | AUC | 1 | 1 | 1 | 1 | 1 | 1 | 1 | 1 | 1 | 0.063 |

**Table 2.** Machine learning results of the top 77, 70, 60, 50, 40, 30, 20 significantly expressed genes, and randomly selected 40 genes in NATs.

|  |  |  |  |  |  |  |  |  |  |  |
| --- | --- | --- | --- | --- | --- | --- | --- | --- | --- | --- |
| Model | Metric | 77 genes | 70 genes | 60 genes | 50 genes | 40 genes | 30 genes | 20 genes | Random 40 genes | Standard Deviation |
| Random Forest | Accuracy | 0.8 | 0.933 | 0.933 | 0.933 | 0.933 | 0.867 | 0.933 | 0.867 | 0.293 |
|  | Precision | 0.75 | 0.9 | 0.9 | 0.9 | 0.9 | 0.818 | 0.9 | 0.818 | 0.507 |
|  | Sensitivity | 1 | 1 | 1 | 1 | 1 | 1 | 1 | 1 | 0.507 |
|  | Specificity | 0.5 | 0.833 | 0.833 | 0.833 | 0.833 | 0.667 | 0.833 | 0.667 | 0.473 |
|  | F1 Score | 0.857 | 0.947 | 0.947 | 0.947 | 0.947 | 0.9 | 0.947 | 0.9 | 0.507 |
|  | MCC | 0.612 | 0.866 | 0.866 | 0.866 | 0.866 | 0.739 | 0.866 | 0.739 | 0 |
|  | AUC | 0.981 | 0.981 | 1 | 1 | 1 | 1 | 1 | 0.907 | 0.009 |
| Logistic Regression | Accuracy | 0.933 | 0.933 | 0.933 | 0.933 | 0.933 | 0.933 | 0.933 | 0.6 | 0.258 |
|  | Precision | 0.9 | 0.9 | 0.9 | 0.9 | 0.9 | 0.9 | 0.9 | 0.667 | 0.507 |
|  | Sensitivity | 1 | 1 | 1 | 1 | 1 | 1 | 1 | 0.667 | 0.507 |
|  | Specificity | 0.833 | 0.833 | 0.833 | 0.833 | 0.833 | 0.833 | 0.833 | 0.5 | 0.487 |
|  | F1 Score | 0.947 | 0.947 | 0.947 | 0.947 | 0.947 | 0.947 | 0.947 | 0.667 | 0.507 |
|  | MCC | 0.866 | 0.866 | 0.866 | 0.866 | 0.866 | 0.866 | 0.866 | 0.167 | 0 |
|  | AUC | 1 | 1 | 0.981 | 0.981 | 1 | 0.981 | 0.981 | 0.759 | 0.032 |
| XGBoost | Accuracy | 0.867 | 0.867 | 0.867 | 0.867 | 0.867 | 0.867 | 0.867 | 0.867 | 0.338 |
|  | Precision | 0.9 | 0.9 | 0.9 | 0.9 | 0.9 | 0.9 | 0.9 | 0.818 | 0.507 |
|  | Sensitivity | 1 | 1 | 1 | 1 | 1 | 1 | 1 | 1 | 0.507 |
|  | Specificity | 0.833 | 0.833 | 0.833 | 0.833 | 0.833 | 0.833 | 0.833 | 0.667 | 0.462 |
|  | F1 Score | 0.947 | 0.947 | 0.947 | 0.947 | 0.947 | 0.947 | 0.947 | 0.9 | 0.507 |
|  | MCC | 0.866 | 0.866 | 0.866 | 0.866 | 0.739 | 0.739 | 0.866 | 0.739 | 0 |
|  | AUC | 1 | 1 | 1 | 1 | 1 | 1 | 1 | 1 | 0 |

To establish a robust baseline, 40 genes were randomly selected from the full set of mitochondrial sense and NATs transcripts and used as machine learning features [Supplementary Data-4]. This control experiment was designed to validate the predictive strength of statistically derived features. The outcome clearly demonstrated that without prior feature selection, the machine learning models failed to yield reliable predictive performance highlighting the critical role of informed, data-driven feature selection in enhancing model accuracy and biological interpretability.

### Evaluation of Predictive Performance by Permutation Testing

Across all tested models and gene sets, the permutation testing consistently yielded low p-values (<0.05) for Accuracy, F1 score, MCC, and AUC, indicating that the observed performance was highly unlikely to arise by chance (Table 3, 4). For the sense gene datasets, the top-performing 80 gene set with Random Forest attained an observed F1 score of 0.941, with the corresponding null distribution producing a markedly lower mean score (p=0.005). Logistic Regression and XGBoost also showed statistically significant enrichment despite comparatively weaker absolute performance (F1=0.896, p=0.005). Likewise, for the NATs 70 gene set, Random Forest achieved an observed F1 score of 0.947, while the mean permuted F1 score was substantially lower, yielding an empirical p-value of 0.005. Similar results were obtained for Logistic Regression and XGBoost. The consistent statistical separation between observed and null distributions across multiple independent models strengthens confidence that the classifiers are capturing biologically relevant signal rather than random structure.

**Table 3.** Permutation results of the top 92, 80, 70, 60, 50, 40, 30, 20 significantly expressed genes in Sense.

| Model | Metric | 92 genes | 80 genes | 70 genes | 60 genes | 50 genes | 40 genes | 30 genes | 20 genes |
| --- | --- | --- | --- | --- | --- | --- | --- | --- | --- |
| Random Forest | Accuracy | 0.01 | 0.01 | 0.005 | 0.005 | 0.005 | 0.005 | 0.005 | 0.01 |
|  | Precision | 0.005 | 0.01 | 0.005 | 0.005 | 0.005 | 0.005 | 0.005 | 0.0149 |
|  | Recall | 0.0597 | 0.0199 | 0.005 | 0.01 | 0.01 | 0.005 | 0.01 | 0.0149 |
|  | MCC | 0.01 | 0.01 | 0.005 | 0.005 | 0.005 | 0.005 | 0.005 | 0.01 |
|  | F1 Score | 0.01 | 0.01 | 0.005 | 0.005 | 0.005 | 0.005 | 0.005 | 0.01 |
|  | AUC | 0.005 | 0.005 | 0.005 | 0.005 | 0.005 | 0.005 | 0.005 | 0.01 |
| Logistic Regression | Accuracy | 0.005 | 0.01 | 0.005 | 0.01 | 0.005 | 0.005 | 0.005 | 0.005 |
|  | Precision | 0.005 | 0.01 | 0.005 | 0.01 | 0.01 | 0.005 | 0.005 | 0.005 |
|  | Recall | 0.01 | 0.01 | 0.0249 | 0.0149 | 0.005 | 0.01 | 0.01 | 0.005 |
|  | MCC | 0.005 | 0.01 | 0.005 | 0.01 | 0.005 | 0.005 | 0.005 | 0.005 |
|  | F1 Score | 0.005 | 0.01 | 0.005 | 0.01 | 0.005 | 0.005 | 0.005 | 0.005 |
|  | AUC | 0.005 | 0.005 | 0.01 | 0.01 | 0.005 | 0.005 | 0.005 | 0.005 |
| XGBoost | Accuracy | 0.1095 | 0.1045 | 0.1244 | 0.0746 | 0.1045 | 0.1045 | 0.0945 | 0.1045 |
|  | Precision | 0.199 | 0.1891 | 0.194 | 0.1443 | 0.1592 | 0.199 | 0.1692 | 0.1642 |
|  | Recall | 0.0249 | 0.0149 | 0.0199 | 0.0149 | 0.0199 | 0.01 | 0.0199 | 0.0249 |
|  | MCC | 0.0547 | 0.0448 | 0.0796 | 0.0299 | 0.0647 | 0.0547 | 0.0547 | 0.0647 |
|  | F1 Score | 0.0547 | 0.0448 | 0.0796 | 0.0249 | 0.0448 | 0.0348 | 0.0498 | 0.0647 |
|  | AUC | 0.005 | 0.005 | 0.01 | 0.005 | 0.005 | 0.005 | 0.005 | 0.005 |

**Table 4.** Permutation results of the top 77, 70, 60, 50, 40, 30, 20 significantly expressed genes in NATs.

| Model | Metric | 77 genes | 70 genes | 60 genes | 50 genes | 40 genes | 30 genes | 20 genes |
| --- | --- | --- | --- | --- | --- | --- | --- | --- |
| Random Forest | Accuracy | 0.0199 | 0.005 | 0.005 | 0.005 | 0.005 | 0.01 | 0.005 |
|  | Precision | 0.0846 | 0.005 | 0.005 | 0.005 | 0.005 | 0.0149 | 0.005 |
|  | Recall | 0.0199 | 0.0149 | 0.0249 | 0.0149 | 0.0149 | 0.01 | 0.0149 |
|  | MCC | 0.01 | 0.005 | 0.005 | 0.005 | 0.005 | 0.005 | 0.005 |
|  | F1 Score | 0.01 | 0.005 | 0.005 | 0.005 | 0.005 | 0.005 | 0.005 |
|  | AUC | 0.005 | 0.005 | 0.005 | 0.005 | 0.005 | 0.005 | 0.005 |
| Logistic Regression | Accuracy | 0.01 | 0.0149 | 0.01 | 0.005 | 0.0149 | 0.005 | 0.01 |
|  | Precision | 0.01 | 0.0149 | 0.01 | 0.005 | 0.0149 | 0.005 | 0.01 |
|  | Recall | 0.0149 | 0.01 | 0.005 | 0.0149 | 0.0149 | 0.0199 | 0.0149 |
|  | MCC | 0.01 | 0.0149 | 0.01 | 0.005 | 0.0149 | 0.005 | 0.01 |
|  | F1 Score | 0.01 | 0.01 | 0.005 | 0.005 | 0.0149 | 0.005 | 0.01 |
|  | AUC | 0.005 | 0.005 | 0.005 | 0.005 | 0.01 | 0.005 | 0.0149 |
| XGBoost | Accuracy | 0.0448 | 0.0348 | 0.01 | 0.0149 | 0.0149 | 0.0149 | 0.01 |
|  | Precision | 0.0547 | 0.0348 | 0.0149 | 0.0149 | 0.0149 | 0.0149 | 0.01 |
|  | Recall | 0.0299 | 0.0149 | 0.005 | 0.01 | 0.0149 | 0.01 | 0.0199 |
|  | MCC | 0.0448 | 0.0348 | 0.01 | 0.0149 | 0.0149 | 0.0149 | 0.01 |
|  | F1 Score | 0.0249 | 0.0149 | 0.005 | 0.01 | 0.005 | 0.005 | 0.01 |
|  | AUC | 0.0149 | 0.0149 | 0.005 | 0.01 | 0.005 | 0.005 | 0.01 |

### Gene ontology (GO) enrichment analysis of Top-Performing Predictive Gene Set

Gene Ontology (GO) analysis was employed to characterize the biological attributes of the genes. For sense transcripts, the top 80 genes were stratified into 57 upregulated and 23 downregulated genes, and GO enrichment was performed on each group independently to capture biological processes specifically associated with up and downregulation (Supplementary Data-5). The GO enrichment analysis revealed significant functional clustering exclusively within the upregulated gene set, while the 23 downregulated genes showed no statistically significant enrichment. The upregulated genes demonstrated a highly coordinated mitochondrial gene regulatory program centred on interconnected metabolic networks essential for *P. vivax* adaptation within host erythrocytes. The most prominent enrichments included amino acid catabolism (GO:0006546, GO:0009071), protein synthesis machinery (GO:0006412), and thiamine diphosphate metabolism (GO:0042357). The simultaneous activation of the glycine cleavage system, thiamine biosynthesis, and mitochondrial translation machinery establishes a metabolic interdependency network wherein glycine catabolism generates one-carbon units for nucleotide biosynthesis while requiring thiamine diphosphate as an essential cofactor for TCA cycle enzymes. Additionally, endocytic pathways (clathrin-dependent and receptor-mediated endocytosis were significantly enriched, indicating coordinated nutrient acquisition mechanisms. This upregulation-dominated regulatory pattern indicates that the parasite’s adaptive response involves selective activation of critical survival pathways, representing a metabolic focusing strategy that concentrates cellular resources on mitochondrial energy production, protein synthesis, and host cell exploitation.

In NATs, the top 70 predictive genes were stratified into 23 upregulated and 47 downregulated genes (Supplementary Data-5). The enrichment analysis of the upregulated transcripts revealed a tightly coordinated program of mitochondrial support functions. The lone detection of prolyl□tRNA aminoacylation (GO:0006433) underscores changes in charging of mitochondrial tRNAs, which can perturb organellar translation. Concurrently, glycine decarboxylation via the glycine cleavage system (GO:0019464) and thiamine diphosphate metabolic and biosynthetic processes (GO:0042357 & GO:0009229) establish a critical one□carbon and cofactor supply network that fuels TCA cycle dehydrogenases and mitochondrial gene expression. Enhanced translational termination (GO:0006415) and complex disassembly processes (GO:0043624 & GO:0032984) indicate active turnover of mitochondrial ribonucleoprotein assemblies. In contrast to the highly focused upregulation program, the downregulated NATs genes were enriched in distinct biological processes primarily linked to mitochondrial function, RNA processing, and protein regulation. Notably, heme metabolic and biosynthetic processes (GO:0046160 & GO:0006784) and respiratory chain complex IV assembly (GO:0008535) were significantly downregulated, indicating a deliberate down-tuning of cytochrome□c oxidase biogenesis. Concurrent downregulation of dolichol and polyprenol biosynthetic/metabolic processes (GO:0019408, GO:0016094 & GO:0016539) suggests reduced N□glycosylation capacity, which may modulate protein trafficking to the mitochondrion. Additionally, suppression of mRNA 3□end processing, polyadenylation, and cleavage (GO:0031124, GO:0006378 & GO:0006379) points to global attenuation of transcript maturation, with modest decrease in translation (GO:0006412), this suggests a strategic conservation of resources by limiting protein synthesis and mitochondrial electron transport chain assembly under specific physiological conditions.

### Pathway enrichment analysis of Top-Performing Predictive Gene Set

In addition to the Gene Ontology, the pathway analysis was also performed to assess the functional relevance, the top-performing genes were systematically mapped to KEGG [40] and MetaCyc [41] pathways to identify activated and suppressed processes across classes. Pathway analysis of up and downregulated genes in both sense and NATs was performed using PlasmoDB [26]. For sense transcripts, 57 upregulated and 23 downregulated genes were mapped to 6 MetaCyc and 1 KEGG pathway while for NATs, 23 upregulated and 47 downregulated genes were mapped to 15 MetaCyc and 5 KEGG pathways (Supplementary Data-6).

Among the upregulated sense transcripts, significant enrichment was observed in thiamine-tRNA regulatory network essential for mitochondrial bioenergetics. The most significant enrichment was thiamine diphosphate salvage III accompanied by multiple thiamine biosynthetic pathways and tRNA splicing. This coordinated upregulation establishes dual thiamine acquisition strategies (salvage and de *novo* biosynthesis) that ensure continuous supply of thiamine diphosphate (ThDP) cofactor for critical mitochondrial dehydrogenases in the TCA cycle. In parallel, enrichment of the tRNA splicing pathway points to elevated processing and maturation of mitochondrial and cytosolic tRNAs, reinforcing translational capacity and mitochondrial gene expression. Together, these changes, highlight an upregulation of cofactor metabolism and translational machinery, indicative of heightened mitochondrial activity.

In contrast, among the downregulated sense transcripts, the only significantly enriched pathway was aminobenzoate degradation. Aminobenzoate is a precursor in folate biosynthesis, which contributes one-carbon units critical for nucleotide synthesis and mitochondrial redox balance. Its suppression suggests a reduced flux toward folate turnover and one-carbon metabolism, potentially restricting cofactor recycling and limiting mitochondrial biosynthetic capacity. This downregulation may reflect an adaptive strategy of the parasite to selectively repress certain mitochondrial metabolic routes while prioritizing cofactor synthesis and translational processes.

In case of NATs, pathway enrichment analysis of upregulated NATs revealed a strong convergence on thiamine diphosphate (TPP) metabolism, with multiple salvage and biosynthetic routes. As TPP is a key cofactor for mitochondrial enzymes driving the TCA cycle, this suggests modulation of cofactor availability to sustain energy metabolism. Additional enrichment in L-cysteine biosynthesis and xenobiotic degradation pathways points to improved redox buffering and metabolic adaptability. Collectively, upregulated NATs appear to reinforce mitochondrial function by boosting cofactor supply, redox stability, and nutrient flexibility, making NATs may act as critical modulators of mitochondrial gene regulation and energy balance in *Plasmodium*.

Interestingly, downregulated NATs highlighted a coordinated suppression of mitochondrial energy metabolism. Key enzymes involved in pyruvate decarboxylation to acetyl-CoA, valine/leucine/isoleucine degradation and propanoate metabolism were significantly downregulated, pointing to reduced anaplerotic input into the TCA cycle. Consistently, both the citrate cycle and pyruvate metabolism were also enriched among suppressed genes, suggesting diminished flux through central carbon metabolism. Parallel downregulation of glycolysis/gluconeogenesis further underscores a global restriction of energy-yielding pathways. In addition, suppression of dolichol and polyisoprenoid biosynthesis pathways together with aminoacyl-tRNA biosynthesis, points toward impaired protein glycosylation and translational capacity, processes tightly coupled with mitochondrial activity.

Together, while sense transcripts mapped largely to thiamine diphosphate salvage and amino acid metabolism, NATs displayed a more profound regulatory footprint by modulating both anabolic (cofactor biosynthesis, cysteine metabolism) and catabolic (pyruvate decarboxylation, TCA cycle, glycolysis) pathways. This dual influence underscores the pivotal role of NATs in fine-tuning a regulatory influence on mitochondrial function in *Plasmodium vivax*, balancing energy generation and metabolic flexibility that may critically determine parasite survival and pathogenicity

### Feature Importance analysis

To identify the key contributors among the top performing genes, SHAP analysis was applied to the best-fitting Random Forest model to interpret feature importance. The top-ranking genes in sense and NATs transcripts (Fig. 2A-2B) demonstrated strong predictive relevance, from the overall 80 genes selected for sense and 70 genes for NATs, providing the most reliable prediction performance across all algorithms tested.

**Figure 2A.**
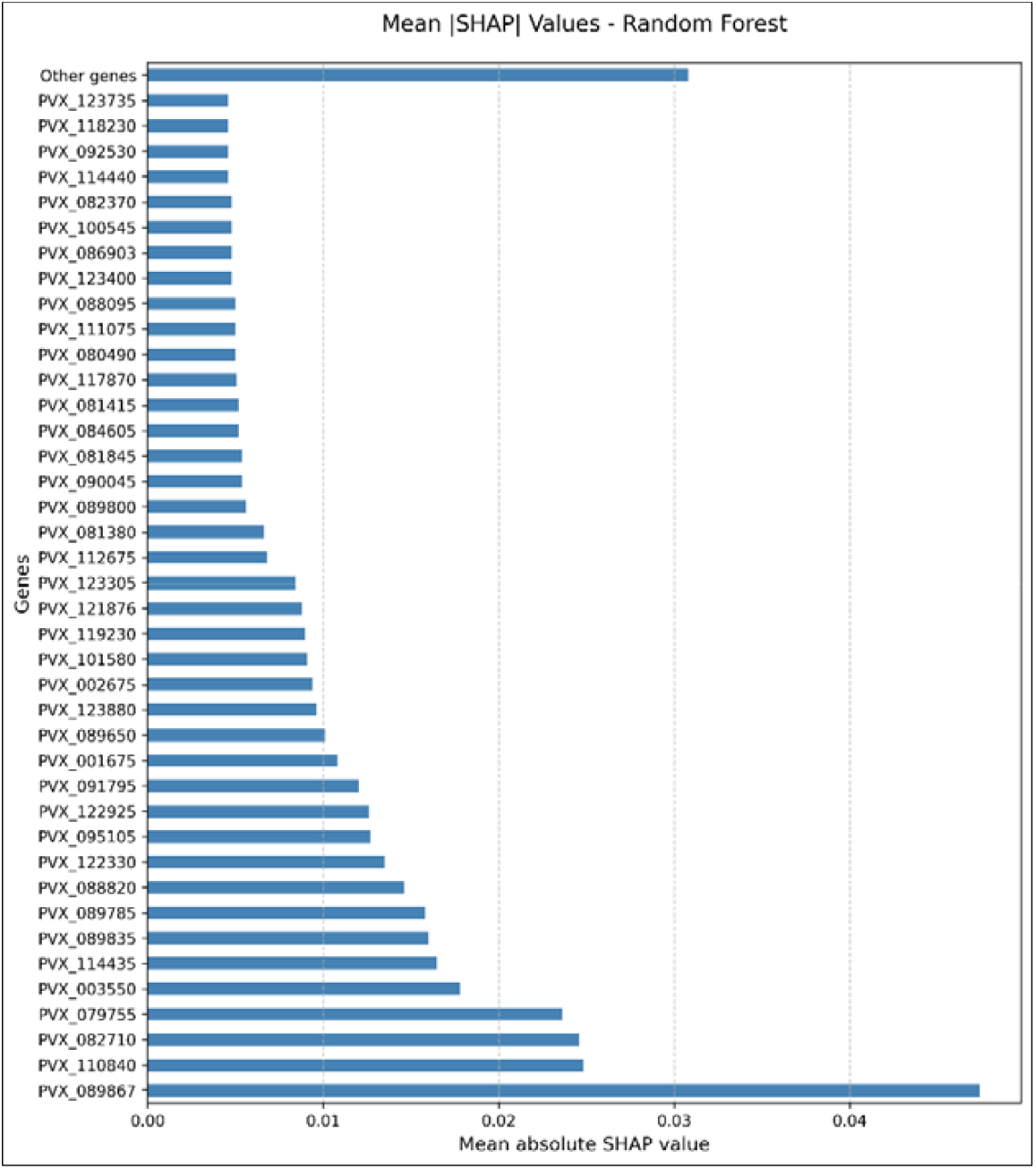
SHAP analysis highlighting key feature importance genes identified by the Random Forest model for sense transcripts

**Figure 2B.**
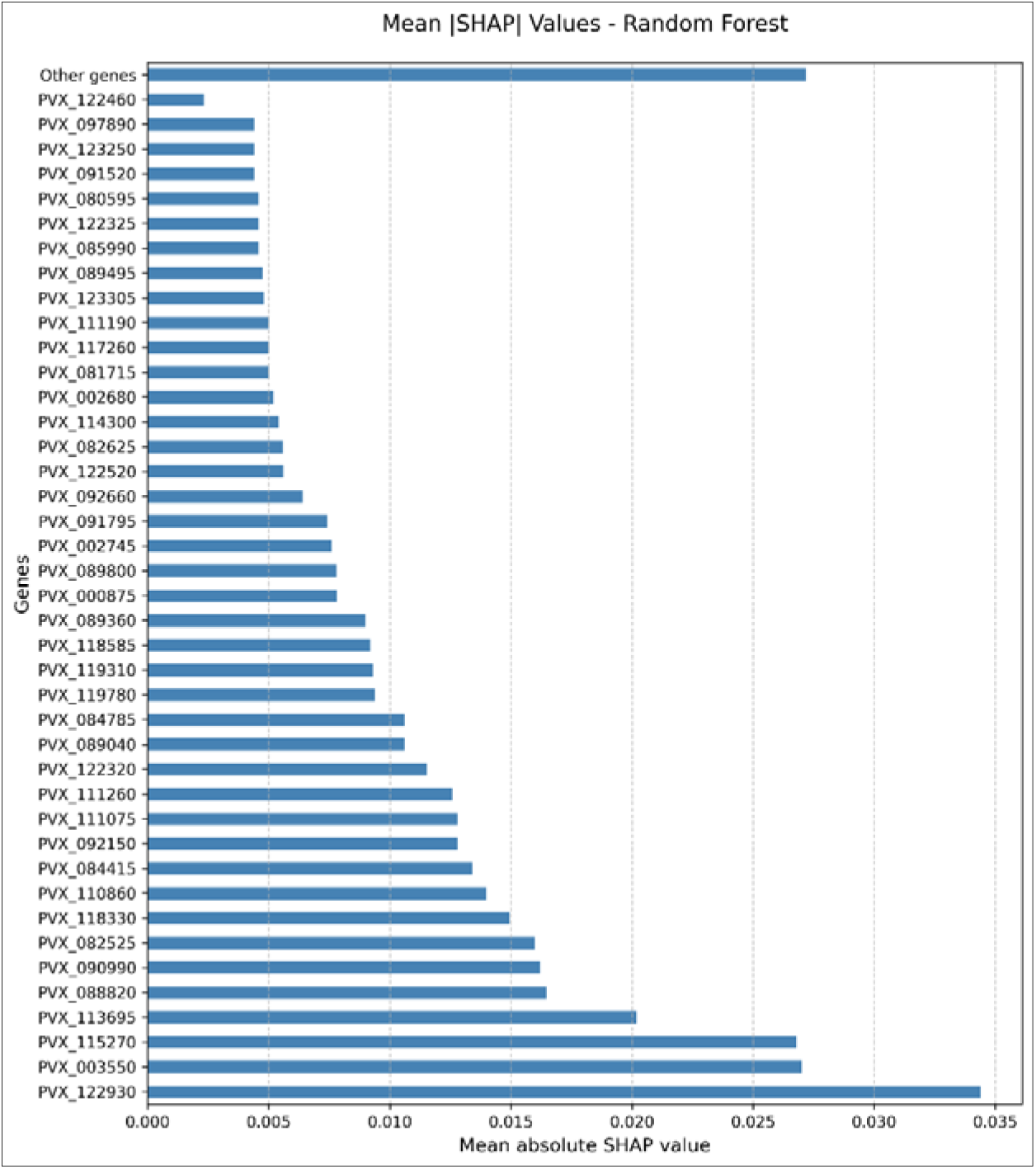
SHAP analysis highlighting key feature importance genes identified by the Random Forest model for NATs

Genes with SHAP values were considered to have a greater influence on the model’s predictions and are potential candidates for involvement in mitochondrial regulatory processes. Notably, genes prioritized by feature importance overlapped with enriched pathways related to polyisoprenoid pathway, amino acid catabolism, mitochondrial translation, and central carbon metabolism, thereby linking computational prioritization with biologically meaningful processes.

To further investigate the directionality of feature contributions, SHAP scatter density plots were generated separately for sense and NATs (Fig. 3A-3B). In these plots, each dot corresponds to a sample, with the x-axis representing the SHAP value (impact on model prediction) and the color indicating relative expression (red = high expression, blue = low expression).

**Fig. 3A.**
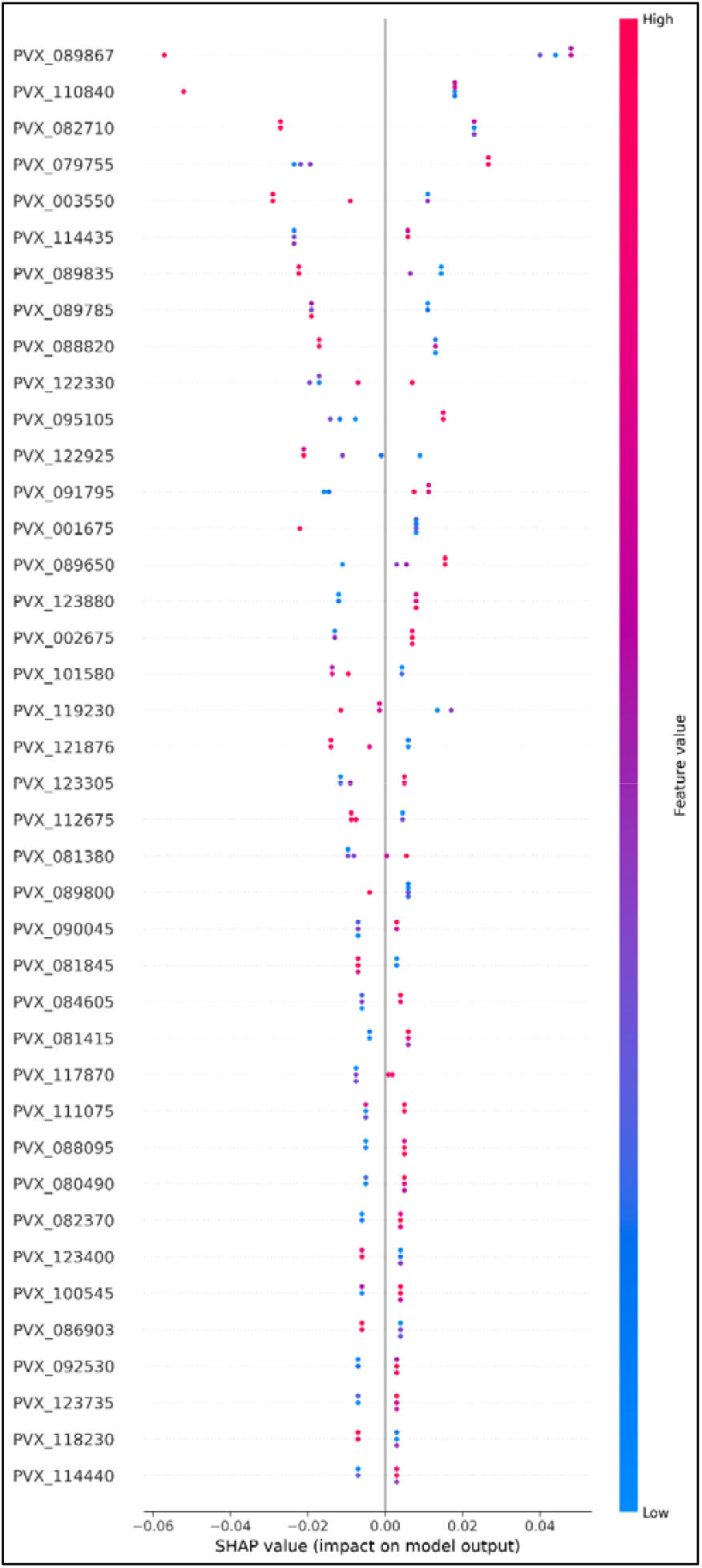
Scatter density SHAP analysis for key feature importance genes identified by the Random Forest model for Sense Dot colour indicates feature value (red = high, blue = low), and x-axis shows each gene’s impact on prediction.

**Fig. 3B.**
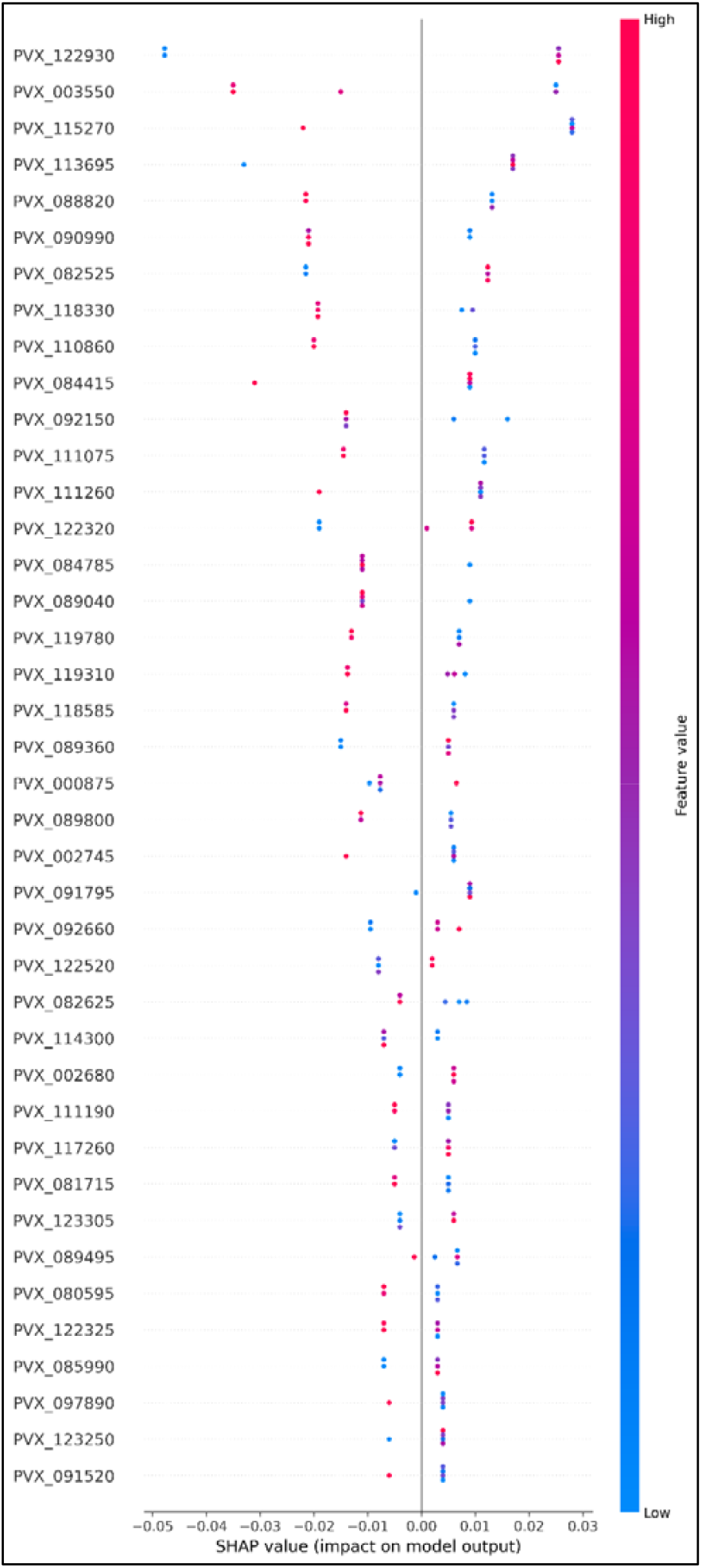
Scatter density SHAP analysis for key feature importance genes identified by the Random Forest model for NATs Dot colour indicates feature value (red = high, blue = low), and x-axis shows each gene’s impact on prediction.

For sense, transcripts such as PVX_095105, PVX_001675, PVX_089650 showed predominantly positive SHAP values, indicating a strong and consistent contribution to the model. Other genes, such as PVX_002675, PVX_084605, PVX_117870, exhibited mixed SHAP values (both positive and negative), suggesting that their regulatory role may be context-dependent or influenced by differential expression across clinical isolates. However, these features were not directly represented in the mitochondrial pathways identified through enrichment analysis.

However, in the case of NATs, transcripts including PVX_089040, PVX_082525, PVX_084785, PVX_090990 and others were associated with consistently positive SHAP contributions, highlighting few of them as potential candidates for mitochondrial regulation. Among transcripts showing mixed SHAP contributions, only PVX_119310 mapped to mitochondrial pathways, suggesting that their contribution to mitochondrial regulation may be context-dependent or vary across clinical isolates.

Collectively, these scatter density plots complement the feature ranking by providing critical insights into the directionality of gene contributions, revealing that transcript abundance in both sense and NATs, with NATs demonstrating a stronger regulatory footprint.

Importantly, SHAP helped us to distinguish which genes were truly influential in driving the model’s predictions, rather than being statistical noise. In this case, SHAP facilitated the prioritisation of candidate genes that may play regulatory roles in mitochondrial pathways, especially among NATs, thereby generating biologically meaningful insights that could guide future functional validation and mechanistic studies.

## Discussion

The study delivers an integrative framework combining transcriptomic profiling and machine learning to unravel mitochondrial gene regulation in severe *P. vivax* malaria, with a particular emphasis on the biological significance of NATs. These findings present a paradigm-shifting view into how mitochondrial dynamics are orchestrated in the parasite and highlight the critical, multifaceted roles of both sense and NATs transcripts.

Our study incorporated transcriptomic data to identify key genes of interest, which were subsequently employed as training sets for model verification. The underlying rationale was that if the model achieved robust predictive accuracy based on these features, it would not only provide predictive power but also strengthen the biological plausibility of the findings. In practice, ML facilitated dimensionality reduction by refining a broad set of statistically significant transcripts into a smaller, reproducible subset of genes. Thus, ML served a major role for enhancing predictive validation thereby reducing potential false positives and uncovering mechanistic leads for future investigation. This integrative framework provides a more interpretable approach for extracting biologically meaningful insights from complex transcriptomic datasets.

The predominance of upregulated sense mitochondrial transcripts in complicated malaria reflects a possible environment driven modulation of mitochondrial metabolic programs under host-imposed stress. Enrichment of pathways mainly in the sense transcripts were amino acid catabolism, mitochondrial translation machinery, and thiamine diphosphate metabolism strongly suggests a strategic metabolic focusing wherein *P. vivax* optimizes energy production and biosynthetic processes critical for survival [14, 41, 42]. The glycine cleavage system and thiamine biosynthesis form an interdependent metabolic network providing crucial one-carbon units and cofactors necessary for mitochondrial enzymatic reactions, highlighting a tightly coordinated metabolic adaptation [39, 40]. Such reprogramming likely supports increased mitochondrial bioenergetic demands during the erythrocytic stage, consistent with reported roles of the *Plasmodium* mitochondrion in maintaining parasite viability [13, 25]. In addition to this, the downregulation of sense transcripts hints towards a nuanced reallocation of mitochondrial resources favouring energy and cofactor production over certain anabolic pathways, likely as a cost-saving measure under infection-induced physiological stress. This aligns with the concept of metabolic focusing where parasites selectively activate essential pathways while suppressing non-critical routes to optimize survival against host immune responses [43], or severe disease pressure.

The landscape of NATs presents a more intricate regulatory signature. The upregulated NATs are enriched in processes essential for mitochondrial translational capacity and cofactor biosynthesis, including prolyl-tRNA aminoacylation, thiamine diphosphate metabolism, glycine decarboxylation, and translational termination [23, 24, 40]. These functions underscore modulation of the cofactor supply, which are prerequisites for efficient translation of mitochondrial-encoded proteins and metabolic activity [14]. In contrast, pronounced downregulation of NATs involved in pathways such as pyruvate metabolism, TCA cycle, glycolysis, and respiratory chain complex IV assembly, points toward a modulation of energy-generating processes. This likely represents a strategic metabolic trade-off facilitating conservation of resources and limiting mitochondrial activity when faced with host immune pressure [28, 42]. Additionally, reduced expression in dolichol and polyisoprenoid biosynthesis may modulate protein glycosylation and mitochondrial protein import, processes critical for organellar integrity and function [25]. The concomitant downregulation of RNA processing and polyadenylation pathways further suggests a fine-tuned control of translation, reinforcing the view of NATs as dynamic regulators in response to physiological states [11, 44].

Interestingly, analysis of concordant expression of the NATs and sense transcripts revealed striking findings. Notably, PVX_099630 (thiamine pyrophosphokinase, putative), a pivotal enzyme in the thiamine diphosphate salvage III pathway, exhibited simultaneous upregulation of both sense transcripts and NATs, underscoring a potential regulatory coordination. This could mechanistically indicate that the NATs act as molecular buffers that prevent protein levels from rising proportionally to mRNA levels. The coordinate upregulation during severe malaria suggests this as a possible stress adaptation mechanism. The parasite enhances metabolic capacity for survival while preventing potentially lethal metabolic imbalances could compromise cellular integrity. A similar phenomenon is observed in case of PVX_092250 (glycine cleavage system H protein, putative), which has shown significant association with GO terms suggesting different catabolic processes (Glycine catabolic process, serine family amino acid catabolic process) amongst others, where both the sense transcripts and NATs are upregulated in severe malaria. The glycine cleavage system and amino acid catabolic pathways identified are metabolically powerful. Uncontrolled activity could lead to excessive NADH production disrupting cellular redox balance or amino acid depletion respectively. This has the potential of affecting other cellular processes [45]. In both the above-mentioned cases, the NATs could be providing a mechanism which would prevent cellular damage from uncontrolled enzyme production during severe malaria and could be part of a sophisticated survival strategy.

The use of LOOCV ensured that each sample was evaluated exactly once, thus preventing performance estimates from depending on a single arbitrary split. Nevertheless, given the limited dataset size, variance across folds remained high, as reflected in the observed standard deviations, an expected limitation of small clinical cohorts. This design helps detect overfitting to specific individuals by repeatedly testing generalization across all samples. While LOOCV reduces bias in small datasets, it does not resolve high variance. To strengthen confidence in the predictive performance, we complemented LOOCV with rigorous permutation testing, which established that the observed accuracy was significantly greater than chance expectations, addressing concerns about overfitting in small clinical cohorts [37]. This addresses the concern that randomly selected genes or latent co-expression modules might drive spurious results. Importantly, the low p-values across most metrics and gene sets confirm that the predictive performance reported here reflects true signal in the gene expression data rather than overfitting to noise. Despite these constraints, the use of machine learning classifiers with LOOCV and SHAP interpretation delivers validation of transcriptomic signatures, confirming the predictive power and biological significance of mitochondrial sense and NATs transcripts [34,35,38]. The consistently high F1 scores, MCC, and AUC metrics highlight the reproducibility and reliability of identified features. Notably, SHAP values emphasize NAT-associated transcripts as having stronger and more stable contributions, underscoring their integral role in mitochondrial gene regulation [46].

This study elucidates NATs as master regulators of mitochondrial function in *P. vivax*, capable of orchestrating both activation and suppression of critical metabolic and translational pathways. By modulating cofactor biosynthesis, translation, and energy metabolism, NATs afford the parasite metabolic plasticity to adapt to the hostile host environment during severe malaria [47,48]. This refined regulatory control extends beyond mitochondrial gene expression, potentially affecting pathways critical for parasite development and drug resistance [20].

The identification of NATs as key modulators of mitochondrial regulators of both metabolic pathways and translational control suggests sophisticated regulatory networks that could be utilized for drug development. The robust predictive performance of our machine learning models alongside transcriptomic data demonstrates mitochondrial NATs as potential molecular players in clinical applications. Despite the robustness of our integrative approach, certain limitations warrant acknowledgment, first, the clinical cohort size was modest, reflecting the inherent challenges of obtaining severe *P. vivax* samples. Although the use of LOOCV maximized data utilization and reduced bias relative to a single train test split, it cannot fully eliminate the risk of overfitting or guarantee generalizability. In small datasets where the number of features vastly exceeds the number of samples, model performance estimates may exhibit high variance even when average metrics appear robust. However, studies have also documented that train/test split approaches produce robust and unbiased performance estimates regardless of sample size [49].

Accordingly, while the SHAP rankings and predictive accuracies observed here are compelling, they should be regarded as preliminary rather than definitive clinical predictors. The positioning of the *Plasmodium* mitochondrion as a potential target is supported by our findings, yet validation in larger and geographically diverse patient cohorts is essential to confirm these regulatory signatures. Importantly, the unique mitochondrial architecture and NATs mediated regulatory mechanisms uncovered in this study highlight multiple potential avenues ranging from targeting parasite-specific metabolic enzymes to disrupting NATs driven regulation. Future directions should integrate multi-tool mitochondrial annotation, interpretable machine learning models in expanded cohorts and experimental validation of mitochondrial targeted transcripts. Such approaches will not only refine predictive gene signatures and strengthen mechanistic inference but also accelerate the translation of transcriptomic insights into actionable targets for combating malaria.

## Acknowledgments

We thank all the patients and technical workers for participating in and supporting this project. This work is funded by the Indian Council of Medical Research (ICMR) under the Extramural Ad-hoc Scheme (PID: 2019-1121, a multicentric project with AD as PI). We thank Birla Institute of Technology (BITS), Pilani (Pilani campus) and SP Medical College, Bikaner, for providing the necessary facilities required for this work. We also thank Genotypic Technology Pvt. Ltd., Bangalore, India, for the microarray hybridization and consultation regarding data analysis.

## Availability of data and materials

All microarray results have been deposited in the Gene Expression Omnibus database (GSE 300144). All the datasets underlying in this study will be shared on reasonable request to the corresponding author. All data generated are provided in the main text and supplementary data files. All patient meta-data included in this article are included in Table S1. The codes are https://github.com/MPSB-Lab/ML-Microarray.git. Reprint requests should be addressed to AD. All Venn diagram illustrations have been made using Venny [50]. Heatmaps were generated using the Seaborn and Matplotlib libraries, while SHAP plots were created using the SHAP and Matplotlib libraries in python v3.11.13.

## Ethics statement

Ethics Approval - Sample collection was earlier approved by SP Medical College Hospital’s Ethics Committee (No. F(Acad)SPMC/2003/2395). Permission to use these samples for further studies was given through IERB approval No. F29(Acad)SPMC/2020/3151 dated 05.09.2020. Informed consent was obtained from all individual participants included in the study. We sincerely thank all the patients for their voluntary consent and participation in the study.

## Author Contributions

AD conceived the study, acquired funding from ICMR, India, and networked to facilitate the outcome. PR was involved in performing the experimentation, analysis, and data interpretation, designing the figures, and drafting the manuscript under the constant guidance of AD. YG was involved in designing, training, and evaluating the machine learning models used in this study. DKK and SKK supervised sample collection and clinical characterization. All authors read and approved the manuscript.

