## Supplementary material for "Integrative Transcriptomic and Machine Learning Approaches to decipher Mitochondrial Gene Regulation in severe *Plasmodium vivax* Malaria": Supplementary Data-1.docx

| **S.No.** | **Patient Id** | **Age/Sex** | **Clinical Presentation** | **Stage Details** | **Platelet count (cells/mm3)** | **Serum bilirubin**  **(mg/dL)** | **Diagnostic tests for malaria** | | |
| --- | --- | --- | --- | --- | --- | --- | --- | --- | --- |
|  |  |  |  |  |  |  | **PBF** | **RMDTs*** | **PCR** |
| 1 | EA | 50 yr, M | Uncomplicated | 90% Tropho + 10% Schizont | 109000 | 0.9 | + | + | + |
| 2 | EG | 15 yr, M | Uncomplicated | 100% Tropho | 60000 | 2.3 | + | + | + |
| 3 | FW | 60 yr, M | Uncomplicated | 100% Tropho | 84000 | 0.7 | + | + | + |
| 4 | DA | 20 yr, F | Uncomplicated | 80% Tropho + 20% Schizont | 51000 | 2.4 | + | + | + |
| 5 | ED | 22 yr, M | Uncomplicated | 70% Tropho + 30% Schizont | 1.4 lac | 0.7 | + | + | + |
| 6 | CC | 28 yr, F | J, A (Hb. - 6.8) | 60% Tropho + 40% Schizont | 200000 | 3.6 | + | + | + |
| 7 | CQ | 65 yr, M | J | 90% Tropho + 10% Ring | 105000 | 4.27 | + | + | + |
| 8 | ER | 25 yr, F | J | 70% Tropho + 30% Schizont | 112000 | 3.6 | + | + | + |
| 9 | ES | 32 yr, F | J | 90% Tropho + 10% Schizont | 112000 | 4.9 | + | + | + |
| 10 | DE | 26 yr, F | J, A (Hb- 5.4), T | 70% Tropho + 30% Schizont | 55000 | 4.4 | + | + | + |
| 11 | DH | 25 yr, M | J | 90% Tropho + 10% Ring | 1.12 lac | 4.1 | + | + | + |
| 12 | EP | 16 yr, F | J | 90% Tropho + 10% Schizont | 113000 | 3.4 | + | + | + |
| 13 | EY | 30 yr, M | J, T | 100% Tropho | 6000 | 3.3 | + | + | + |
| 14 | EZ | 25 yr, F | J, A (Hb. - 7), T | 60% Tropho + 40% Schizont | 60000 | 5.5 | + | + | + |

**Clinical characteristics of patient samples**

**Table S1: Clinical characteristics of patient samples used in the study**

J- Jaundice, A - Anemia, T -Thrombocytopenia, PBF - Peripheral Blood Film, RMDTs - Rapid Malaria Diagnostic Tests,

PCR - Polymerase Chain Reaction, Hb - haemoglobin (in mg %), ser.bil. Serum bilirubin (in mg %).
